# Identification of Key Differentially Expressed MicroRNAs in Cancer Patients Through Pan-cancer Analysis

**DOI:** 10.1101/388892

**Authors:** Yu Hu, Hayley Dingerdissen, Samir Gupta, Robel Kahsay, Vijay Shanker, Quan Wan, Cheng Yan, Raja Mazumder

**Author notes:** (Hu Y), (Dingerdissen H), (Gupta S), (Kahsay R), (Shanker V), (Wan Q), (Yan C), (Mazumder R). Corresponding author: address: Ross Hall, 2300 Eye Street N.W., Washington, DC 20037.

## Abstract

A number of microRNAs (miRNAs) functioning in gene silencing have been associated with cancer progression. However, common expression patterns of abnormally expressed miRNAs and their potential roles in multiple cancer types have not yet been evaluated. To minimize the difference of patients, we collected miRNA sequencing data of 575 patients with tumor and adjacent non-tumorous tissues from 14 cancer types from The Cancer Genome Atlas (TCGA), and performed differential expression analysis using DESeq2 and edgeR. The results showed that cancer types can be grouped based on the distribution of miRNAs with different expression patterns. We found 81 significantly differentially expressed miRNAs (SDEmiRNAs) unique to one of the 14 cancers may affect patient survival rate, and 21 key SDEmiRNAs (nine overexpressed and 12 under-expressed) associated with at least eight cancers and enriched in more than 60% of patients per cancer, including four newly identified SDEmiRNAs (hsa-mir-4746, hsa-mir-3648, hsa-mir-3687, and hsa-mir-1269a). The downstream effect of these 21 SDEmiRNAs on cellular functions was evaluated through enrichment and pathway analysis of 7,186 protein-coding gene targets from literature mining with known differential expression profiles in cancers. It enables identification of their functional similarity in cell proliferation control across a wide range of cancers and to build common regulatory networks over cancer-related pathways. This is validated by construction of a regulatory network in PI3K pathway. This study provides evidence of the value of further analysis on SDEmiRNAs as potential biomarkers and therapeutic targets for cancer diagnosis and treatment.

## Introduction

MicroRNAs (miRNAs) are small non-coding RNAs with two forms, the premature miRNA (length 50-125 bp) and the processed, mature form (length 18-24 bp)[1]. miRNAs function in gene silencing by binding to mRNAs, causing either mRNA destabilization or inhibition of translation[2]. From the beginning of 21^st^ century, pioneer researchers have increasingly catalogued the close relationship between miRNAs and cancer development. For example, Calin et al.[3] discovered the deletion or down-regulation of miR15/miR16 in chronic lymphocytic leukemia, Takamizawa et al.[4] showed under-expressed hsa-let-7 is related to lower survival rate in lung cancer, Chan et al.[5] knocked down hsa-mir-21 in glioblastoma cells which activated apoptosis, and Li et al.[6] found hsa-mir-10b enabled cell metastasis in breast cancer. These studies not only laid the foundation for a number of studies on the roles of these miRNAs in different cancer types, but also paved the way to the increasing identification of cancer-associated miRNAs[7].

Furthermore, it is believed that there are significantly fewer known human miRNAs (about 1,881 pre-mature and 2,588 mature in miRBase as of 2016[1]) than total mRNAs because each miRNA has multiple target mRNAs. Due to this one-to-many relationship, studying miRNA expression profiles could contribute more efficiently to construction of the molecular regulatory network as a whole, especially in characterizing cancer-associated regulation of molecular pathways or biological processes. miRNAs are classified as tumor suppressors or oncogenes (oncomirs) according to their expression profiles in cancer[8]. In breast cancer, at least 20 oncogenic miRNAs or cluster families have been identified to promote cell proliferation, invasion, migration, or angiogenesis[9]. Cluster families are groups of several miRNAs transcribed together and co-expressed. Similarly, more than 30 tumor suppressive miRNAs or families have been observed to play a role in cell apoptosis and negative control of cell proliferation or migration[9]. In cancer metastasis, these miRNAs can change the expression of genes essential for cellular homeostasis and for robustness of cell fate decisions, often through various signaling pathways[10]. Therefore, to gain a comprehensive view of the cancer landscape, it is imperative to characterize miRNAs on a large scale and integrate evidence that has been observed across multiple cancer types to build more inclusive regulatory networks.

In large-scale studies, roles of miRNAs have been found to be important in cancer diagnosis, prognosis, and therapy for multiple cancer types[10–12]. Each cancer type has a unique miRNA expression signature that can be classified into different prognostic groups[13]. With an increasing number of studies about cancer incidence, there has been a corresponding increase in observations of tumors with different molecular mechanisms even within a single cancer type[14, 15]. Cancer subtypes can also be identified by miRNA expression profiling, as in the case of non-small cell lung cancer where miRNAs have classified three subtypes by different driving factors (ALK, EGFR, and KRAS)[16]. The diagnostic and prognostic roles of miRNAs further indicate their potential as therapeutic targets and as candidates for clinical trials[17]. Recently, miRNAs have been gradually introduced into personalized medicine approaches for cancer or other disease therapies[18, 19]. Based on the apparent importance of miRNAs in cancer, a number of studies have been conducted on miRNA expression profiles in which statistical analysis was used to determine changes between experimental and control samples to characterize cancer-specific miRNAs in one or two cancers. However, pan-cancer analysis of miRNA expression has not been explored to identify similarity and differences across multiple cancers.

With the introduction of newer, more sensitive procedures in library sample preparation, accompanied by decreasing costs of next-generation sequencing (NGS) methods[20], researchers can now acquire a comprehensive miRNA abundance profiling through deep sequencing[21], despite known complications like low abundant miRNAs and miRNA processing complexity. Several normalization methods for miRNA-seq have been widely used in previous studies[22, 23], including edgeR (Trimmed Mean of M values, TMM)[24] and DESeq[25]. In this study, we performed differential expression analysis by both DESeq2[26] and edgeR[24] for paired miRNA-seq data (tumor and surrounding non-tumorous tissues) from The Cancer Genome Atlas (TCGA: http://cancergenome.nih.gov; http://cancergenome.nih.gov; https://portal.gdc.cancer.gov/), following the methods described by Yang S et al[27] and used in the study of Metpally RP et al[20].

Although previous studies have demonstrated critical roles of miRNAs in the prognosis of cancer types and subtypes, our research singled out key miRNAs observed to have significantly differential expression (SDEmiRNAs) for 14 TCGA cancer types independently. The distribution of these SDEmiRNAs with different expression trends enables us to group the 14 cancers and based on different potential developmental mechanisms of the 14 cancers. Individually, comparing the expression profiles of these SDEmiRNAs, we were able to screen out a list of miRNAs which are significantly differential expressed in only one of the 14 cancers. These unique SDEmiRNAs specific to one cancer could be considered as biomarkers of each cancer, and may have influence in patient survival rate, regardless of their driven factors. We also identified a subset of key SDEmiRNAs that are significantly over- or under-expressed with high patient frequencies in at least eight cancer types (over-expression implies functional up-regulation). With the functional and enrichment analyses of their cancer-related target genes and pathways, these SDEmiRNAs might be commonly functional and executive factors promoting multiple cancer development via similar molecular mechanisms. As an example, we constructed common regulatory networks of several SDEmiRNAs in signaling pathways across different cancer types, despite the great difference among their target genes in multiple tissues. Notably, some of the identified SDEmiRNAs are barely studied in cancer development due to their relatively low expression levels but have been found to be significantly differentially expressed in a wide range of cancer types, showing their potential vital roles in cancer development. Thus, this study evaluates the expression of all human miRNAs in 14 cancer types, widens the roles of individual SDEmiRNAs to cancer types where they have not been studied, classifies the functions of SDEmiRNAs in different tissues to small number of molecular mechanisms, and therefore, provides a means for improved discovery of mechanisms of cancer development that could improve downstream clinical studies and trials by prioritizing miRNA and targets for translational study. The results are integrated in BioXpress v3.0 (https://hive.biochemistry.gwu.edu/bioxpress) and open-source to support further analysis.

## Results and Discussion

### Data collection and overall evaluation before differential expression analysis

The TCGA project provides a comprehensive database with a unified data analysis pipeline for collecting and analyzing samples. We were able to collect tumor-only expression data for 1,904 miRNAs in 32 TCGA/ICGC cancer types generated by the HiSeq platform (7,994 TCGA patients and 168 ICGC patients) and 1,668 miRNAs in 12 TCGA cancer types generated by Genome Analyzer (1,439 TCGA patients), from a total of 9,601 patients. Within these data, 22 TCGA cancer types containing paired data (tumor data with matched adjacent non-tumorous data) from the HiSeq platform, we set a threshold of patient counts (greater than 10, covering more than 95% of total patients with paired data), resulting in 14 cancer types to be used in further analysis (BLCA, BRCA, PRAD, STAD, KIRP, KIRC, KICH, LUAD, LUSC, LIHC, THCA, ESCA, UCEC, and HNSC – please see Table 1 for full cancer names). These cancer types involved a total of 575 TCGA-only patients (1,152 paired samples) with an average of 1,393 expressed miRNAs for each cancer.

**Table 1.** Summary of patients and miRNA expression counts, and significantly differentially expressed miRNAs over- and under-expressed counts reported by DESeq2 and edgeR for paired data of 14 TCGA cancer types

| TCGA-Cancer Type | TCGA Cancer Type (full name) | Mapped Cancer Type (DOID) | Patient Counts | Expressed miRNA Counts | miRNA sig over-expressed | miRNA sig under-expressed | Total sig miRNA | 80% patients* |
| --- | --- | --- | --- | --- | --- | --- | --- | --- |
| BLCA | Bladder Urothelial Carcinoma | DOID:11054 / Urinary bladder cancer [UBC] | 19 | 1361 | 18 | 115 | 133 | 72 |
| UCEC | Uterine Corpus Endometrial Carcinoma | DOID:363 / Uterine cancer [Uteca] | 18 | 1370 | 39 | 193 | 232 | 115 |
| LIHC | Liver hepatocellular carcinoma | DOID:3571 / Liver cancer [Livca] | 48 | 1414 | 98 | 70 | 168 | 52 |
| HNSC | Head and Neck squamous cell carcinoma | DOID:11934 / Head and neck cancer [H&NC] | 43 | 1463 | 100 | 152 | 252 | 65 |
| STAD | Stomach adenocarcinoma | DOID:10534 / Stomach cancer [Stoca] | 41 | 1426 | 28 | 126 | 154 | 51 |
| ESCA | Esophageal carcinoma | DOID:5041 / Esophageal cancer [EC] | 12 | 1215 | 4 | 38 | 42 | 24 |
| THCA | Thyroid carcinoma | DOID:1781 / Thyroid cancer [Thyca] | 59 | 1506 | 50 | 116 | 166 | 59 |
| PRAD | Prostate adenocarcinoma | DOID:10283 / Prostate cancer [PCa] | 52 | 1350 | 16 | 64 | 80 | 21 |
| BRCA | Breast invasive carcinoma | DOID:1612 / Breast cancer [BRCA] | 75 | 1445 | 123 | 163 | 286 | 58 |
| LUSC | Lung squamous cell carcinoma | DOID:1324 / Lung cancer [Lunca] & DOID:3907 / Lung squamous cell carcinoma | 44 | 1468 | 89 | 205 | 294 | 74 |
| LUAD | Lung adenocarcinoma | DOID:1324 / Lung cancer [Lunca] & DOID:3910 / Lung adenocarcinoma | 39 | 1403 | 63 | 164 | 227 | 91 |
| KICH | Kidney Chromophobe | DOID:263 / Kidney cancer [Kidca] & DOID:4471 / Chromophobe adenocarcinoma | 24 | 1312 | 148 | 67 | 215 | 119 |
| KIRC | Kidney renal clear cell carcinoma | DOID:263 / Kidney cancer [Kidca] & Kidney renal clear cell carcinoma | 67 | 1363 | 133 | 134 | 267 | 74 |
| KIRP | Kidney renal papillary cell carcinoma | DOID:263 / Kidney cancer [Kidca] & DOID:4465 / Papillary renal cell carcinoma | 34 | 1400 | 135 | 99 | 234 | 85 |

*Note: \*80% patients: counts of miRNAs that are significantly over-expressed (significantly under-expressed) in at least 80% of patients in each cancer type.*

In order to overview the expression profiles of miRNAs in the paired samples from the 14 cancer types, RPM values of each miRNA (pre-calculated by TCGA) in each TCGA patient were first used to calculate regular fold change (FC) values between cancer and corresponding non-tumorous samples. After comparing these FC values, we found that the total number of overexpressed miRNAs is two-fold higher than that of under-expressed miRNAs. However, when considering only those miRNAs with FC > 2 (FC < 0.5 for under-expressed miRNAs), and with relatively high frequencies in patients in at least eight cancers, the trends of the numbers of miRNAs with over- and under-expression are similar to each other across more than 35% of patients in each cancer (Figure 1). This suggests that many over-expressed miRNAs may be effectors driven by other factors and vary from patient to patient.

**Figure 1.**
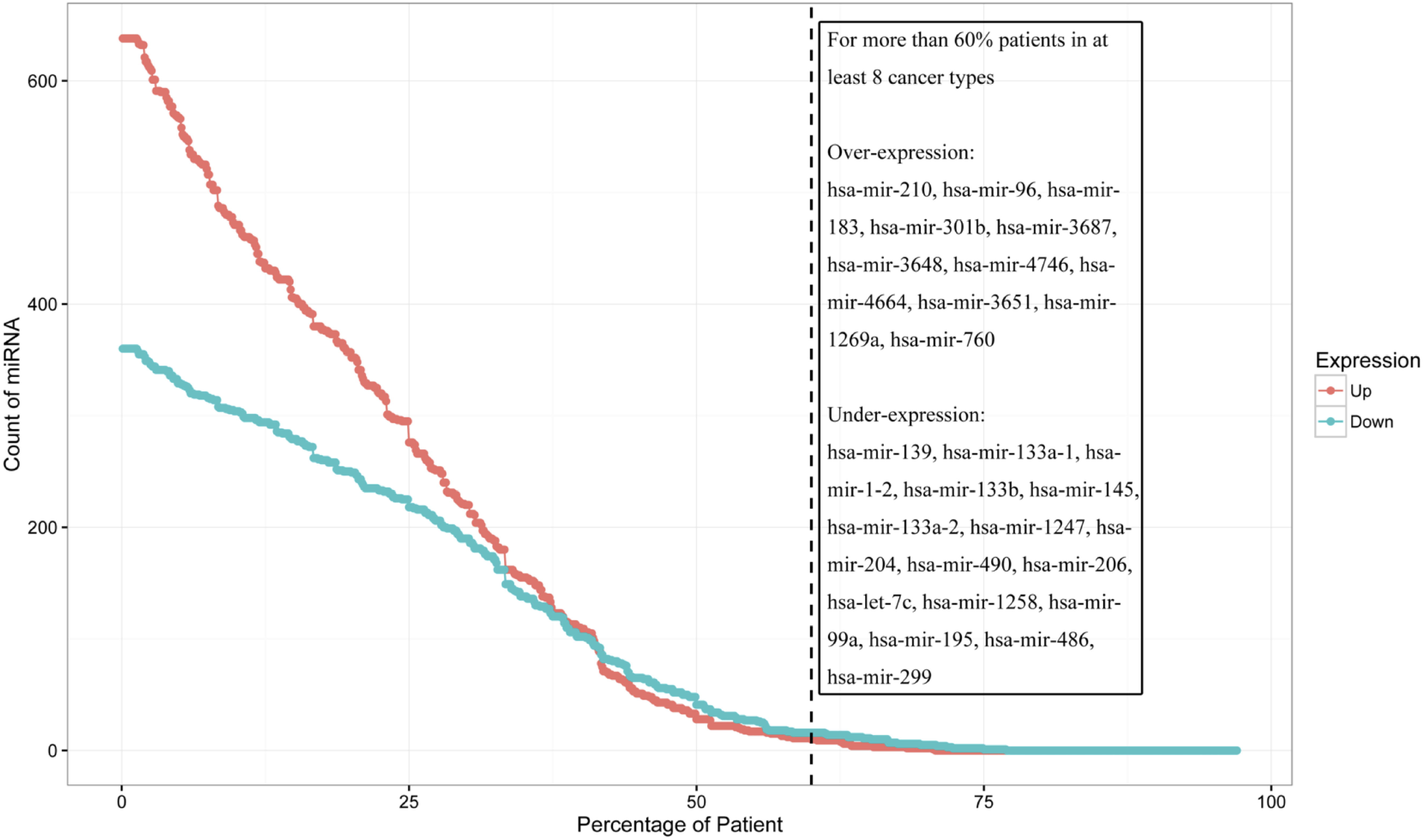
Overview of miRNA counts vs. patient frequency for over- and under-expressed miRNAs with original FC > 2 of FC < 0.5 in at least eight cancer types from paired data. FC is used to distinguish the overall expression trend of miRNAs (by comparing FC values from tumor and non-tumorous samples of the same patient). Each dot on the red line represents the count of over-expressed miRNAs per patient frequency, and dots on the blue line represent counts of under-expressed miRNAs. Across more than 35% of patients, the numbers of miRNAs with over- and under-expression are similar to each other. Those over-expressed miRNAs with low patient frequencies may be effectors of other factors. 11 over-expressed miRNAs and 16 under-expressed miRNAs appear in more than 60% of patients for at least eight cancer types: these miRNAs are listed in the text box. These miRNAs with a high occurrence rate among patients in multiple cancer types suggest their potential roles in the development of these cancers.

This expression distribution also points out 27 miRNAs (11 over-expressed miRNAs and 16 under-expressed miRNAs) in more than 60% of patients (Figure 1) in eight or more cancers. Most of these miRNAs have been previously reported in at least one publication associated with cancer, some of which are well-studied tumor suppressors, including has-mir-133b, and hsa-mir-139, or contribute to cell invasion, such as hsa-mir-301b[28, 29] hsa-mir-1247 was newly found to be a prognostic marker in pancreatic cancer as a suppressor[28], and hsa-mir-1258 targets heparanase (HPSE) to suppress breast and non-small cell lung cancer[30, 31]. hsa-mir-1269a, hsa-mir-3687, and hsa-mir-3648 were also newly identified to be involved in cancer development[32–34], but their molecular mechanisms in cancers need to be researched further. Besides them, two miRNAs reported in 2011[35], hsa-mir-4746 and hsa-mir-4664, which are over-expressed in 11 and eight cancers with average of patient frequencies, 74.42% (average FC = 3.95) and 75.49% (average FC = 8.07), respectively, although their average expression levels are lower than those well-studied miRNAs, which may lead to difficulties in quantification and qualification.

These results validate the reliability and confidence of our data, and suggest the potential similar roles of the subset of 27 miRNAs in multiple cancer development, based on their similar expression profiles. It is also important to note that after statistical analysis in the following sections, 21 out of these 27 miRNAs were found to be significantly differentially expressed (SDEmiRNAs) in the same cancer types.

### miRNA differential expression analysis

#### SDEmiRNAs could be used in distinguishing cancer types

DESeq2[26] and edgeR[24] were used to perform differential expression analysis, applying statistical tests to minimize the potential false positive errors from the differences among individuals and samples. Data from all patients for a single cancer type were analyzed together resulting in reporting average differential expression levels by cancer type. Both DESeq2 and edgeR provide *p* values and adjusted *p* values (or FDR in edgeR). The results of the two tools showed a similar ranking of significantly differentially expressed miRNAs (SDEmiRNAs) when ranked by adjusted *p* values. We found 10 to 15% of miRNAs with significant differential expression in each cancer type reported by both DESeq2 and edgeR (Table 1). As shown in Figure 2A, of all 1,870 miRNAs in our integrated database, the specific distributions of the quantities and types of 656 SDEmiRNAs vary between cancer types, suggesting the existence of unique SDEmiRNA patterns for each of these cancer types. These unique patterns of SDEmiRNAs potentially reduce the amount of miRNAs used for prognosis of cancer types, based on the findings of Lu *et al*[13]. The result of differential expression analysis of each miRNA is available in BioXpress, shown by both figures and table (Figure S1).

**Figure 2.**
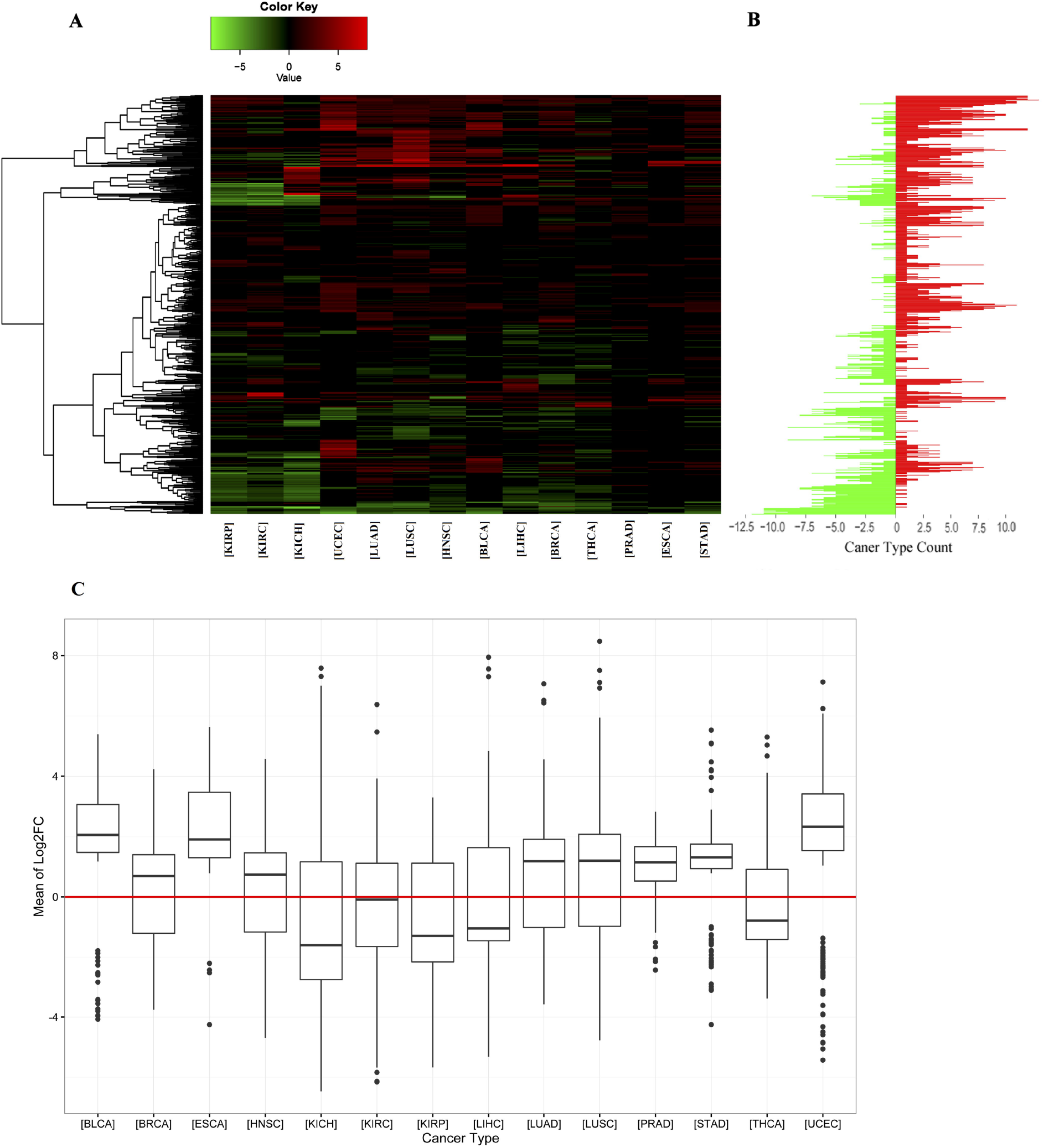
Differential expression analysis of miRNAs and distribution of log2FC values of significantly differentially expressed miRNAs across 14 cancer types. **A.** Heatmap showing significantly over- and under-expressed miRNAs in at least one cancer type - vertical axis is miRNAs. The expression change of miRNA is represented by log2FC values generated by DESeq2 and edgeR. Log2FC values less than zero are considered as under-expressed miRNAs (green), while log2FC greater than zero represents over-expressed miRNAs (red). The miRNA patterns of each cancer type vary from each other. **B.** Counts of cancer types in which each miRNA is over- or under-expressed - vertical axis stands miRNAs (same as Figure 2A). Green bars represent the counts of cancer types in which a miRNA is under-expressed, and red ones are for cancer type counts in which a miRNA is over-expressed. After unsupervised clustering, some miRNAs show the same trend of over-expression or underexpression across most cancers. It suggests that these miRNAs may have similar function across different cancer types. **C.** The vertical axis represents the average of log2FC values generated from DESeq2 and edgeR for each miRNA in each cancer type. For each cancer, all of its miRNAs that are significantly differential expressed are included. The numbers of over- and under-expressed miRNAs per cancer vary from each other, suggesting the up-regulation of oncogenes is more important in the occurrence of some cancers, while the down-regulation of tumor suppression genes is more important in other cancers.

Moreover, similar SDEmiRNA expression patterns are shown in LUAD and LUSC from lung tissue, as well as in KICH, KIRP, and KIRC from kidney tissue, as expected. However, among 387 SDEmiRNAs in either LUAD or LUSC (Figure S2A), 190 have log2FC values greater than 1 (Δlog2FC > 1), 140 (36.2% of 387) of which have the opposite expression trend for both cancers, suggesting they could be key factors underlying the differences between LUAD and LUSC. We also performed a pairwise comparison among KICH, KIRP, and KIRC using the set of 330 SDEmiRNAs with Δlog2FC > 1 in at least one of the three cancer types. It shows that KIRP and KIRC have more similar SDEmiRNA expression patterns than KIRP and KICH, and KICH and KIRC (Figure S2B-D). This suggests that the development of KICH may be mechanistically more distant than the development of KIRC and KIRP.

#### Expression pattern of SDEmiRNAs groups cancer type based on different patterns of mechanisms in different cancer development

We counted the number of SDEmiRNAs that have the same expression trend in at least 80% of patients (by log2FC values) for each cancer type (Table 1, and the corresponding patient frequencies in Figure S3). These results showed different cancer types have different proportions of SDEmiRNAs: some cancers have more over-expressed SDEmiRNAs than underexpressed; some have a balance between over- and under-expressed SDEmiRNAs; and other cancer types have more under-expressed SDEmiRNAs (Figure S3). Similar results are shown when the log2FC values of all SDEmiRNAs were used to display their distributions in all cancer types (Figure 2C).

In four cancer types (KICH, KIRP, LIHC, and THCA), the numbers of under-expressed SDEmiRNAs are much higher than those of over-expressed SDEmiRNAs, while for most of the cancer types (BLCA, ESCA, KIRC, LUAD, LUSC, PRAD, STAD, and UCEC), the median expression values of their SDEmiRNAs are higher than zero. Because the main function of miRNA is to suppress target genes, the significant number of under-expressed miRNAs corresponds to the up-regulation of oncogenes, which may play a more important role in the occurrence and progression of the four cancers characterized predominantly by underexpression of miRNAs. Conversely, the down-regulation of tumor suppression genes may be more important in cancer occurrence and progression for those cancers characterized predominantly by over-expression of miRNAs. Cancers like BRCA, HNSC, and KIRC have an almost symmetric distribution of SDEmiRNAs about log2FC = 0 (Figure 2C), suggesting regulation of both oncogenes and tumor suppression genes could be equally important for disease progression. However, it could also be possible that the similar distribution between over- and under-expressed miRNAs indicates that these miRNAs are equally unimportant to disease and may suggest other mechanisms are primarily at work for these cancer types.

The distribution of SDEmiRNAs is able to group cancer types to three categories based on the potential different mechanisms during cancer initiation and development. It allows high efficiency of further research on exploring mechanisms of cancers.

#### SDEmiRNAs that are unique to one of the 14 cancer types might be driver miRNAs

Although we are mostly interested in SDEmiRNAs implicated in multiple cancers (as discussed in depth below), SDEmiRNAs that only exist in a single cancer type may be considered potential driver miRNAs or biomarkers specific for that cancer type. Therefore, we set up screening conditions to identify those SDEmiRNAs affecting a single cancer type: 1) SDEmiRNA should be reported to be significant by both DESeq2 and edgeR; 2) this SDEmiRNA should not exist in the remaining 13 cancers; 3) the patient frequency of this SDEmiRNA should be more than 50% in this cancer; 4) the log2FC value of this SDEmiRNA should be greater than 1, or smaller than −1.

As shown in Table S1, we identified a total of 81 SDEmiRNAs with significant expression trends in a single cancer type, 57 of which have patient frequencies higher than 60% (Table 2). Most of these SDEmiRNAs have much lower expression in tissues compared with other well-studied miRNAs, and have not been functionally studied in cancer development, especially in their corresponding cancer types. However, this does not negate their potential cancer-related. In order to validate the results, as an example, we performed survival analysis on four SDEmiRNAs unique to BRCA (absolute log2FC values > 1) by using miRpower[36]. Except hsa-mir-329-2 which is not in miRpower, the rest three SDEmiRNAs suggest significantly survival difference (log-rank p value < 0.01) between groups with under-/over-expression of each SDEmiRNA, after optimizing the cutoff of the patients (Figure S4): hsa-mir-4784 (Hazard Ratio = 1.82), hsa-mir-1262 (HR = 0.58), and hsa-mir-320c-1 (HR = 0.68). The difference of survival rates between patients within under- and over-expression were consistent with our results. Even though the input datasets were not exactly matched with this study, it still shows the potential vital roles of these unique SDEmiRNAs in the development of BRCA. Similarly, other SDEmiRNAs may also have the potential as signatures of different cancer types, and are of great importance and prospect to have additional analysis.

**Table 2.**
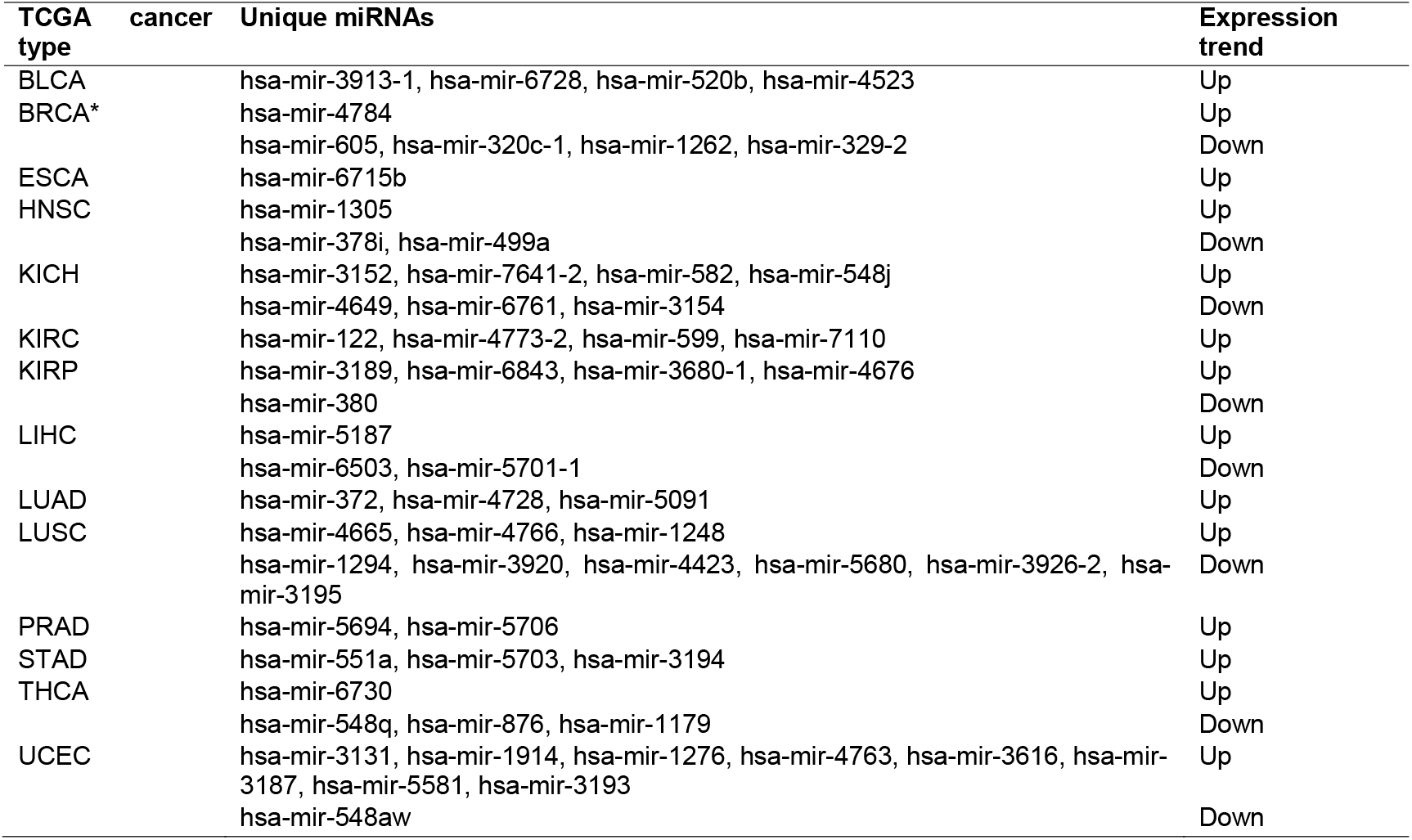
Unique miRNAs with significant differential expression in 60% or more patients of individual cancer

### Pan-cancer analysis of key SDEmiRNAs

As shown in Figure 2B, after unsupervised clustering of all SDEmiRNAs, several SDEmiRNAs follow the same trend of either over-expression or under-expression across most of the cancer types, suggesting that these SDEmiRNAs may function similarly across different cancer types. To further study the key SDEmiRNAs affecting multiple cancers, we identified a subset of 90 miRNAs that have significant differential expression in at least eight cancer types (58% of total cancer types).

#### Selected SDEmiRNAs were validated by experimentally validated databases and text mining tool

Before proceeding with the analysis, we first confirmed these 90 key SDEmiRNAs generated from DESeq2 and edgeR using four experimentally validated databases. Among 2,280 significantly differentially expressed miRNA-cancer correlations, 208 (9.12%) correlations were recorded in the union of the four databases (HMDD, Mir2disease, miRCancer, and miRiaD) to be exclusively over- or under-expressed in certain types of cancer. Furthermore, we randomly picked 24 of 90 SDEmiRNAs (a total of 226 miRNA-cancer correlations) as the input of our text mining tool (DEXTER). After mapping and manual curation, we extracted 157 (69.47%) interactions (Table S2). None of the 90 SDEmiRNAs were found by the databases of DEXTER with opposite expression trends compared to the trends we identified in our study. Based on these observations, we consider our differential expression analysis results to be reliable.

#### Key SDEmiRNAs with at least 60% patient frequencies were considered reliable and may have similar functions across multiple cancers

From the 90 selected SDEmiRNAs, we defined a subset of them based on their expression trend: 47 SDEmiRNAs are all over-expressed and 18 are all under-expressed in eight or more cancer types (a total of 65 SDEmiRNAs with the same significantly differential expression trend in eight or more cancers) (Table S3). Beside 11 SDEmiRNAs, the rest 54 SDEmiRNAs belong to 40 miRNA families, according to MirGeneDB v2.0[37]. Among them, all members of 22 families are included in the list, although most of the families have only one or two members. However, the similar expression trends of these SDEmiRNAs suggest their high consistence of activities and potential functions across 58% of investigated cancers, and therefore, these small families can be classified to larger ones (Table S3). Individually, six newly reported miRNAs were included in the 65 SDEmiRNAs: hsa-mir-3170, hsa-mir-3677, hsa-mir-4326, hsa-mir-4652, hsa-mir-7706, and hsa-mir-105-2 (Table S3). Among them, hsa-mir-4652 is the top one significantly over-expressed miRNAs with largest log2FC value (average log2FC = 5.34, patient frequency larger than 60% in seven cancer types). We found no cancer-related studies that have explored the mechanisms of these six miRNAs in cancers, suggesting their high potential for involvement in cancer development, especially hsa-mir-4652.

Comparing the 65 SDEmiRNAs identified by differential expression analysis to the 27 miRNAs with regular FC > 2 or FC < 0.5 across 60% of patients (in ‘Data collection and evaluation’ section, Figure 1) in at least eight cancer types, we found an overlap of nine overexpressed and 12 under-expressed SDEmiRNAs between the two datasets (a total of 21 miRNAs) (Table 3). This subset not only includes well-studied miRNAs such as hsa-mir-145 and hsa-mir-210, but also newly identified miRNAs such as hsa-mir-4746 (in ‘Data collection and evaluation’ section), hsa-mir-3648, hsa-mir-3687, and hsa-mir-1269a. Two of the overexpressed and five of the under-expressed key miRNAs are in the top five significantly over- and under-expressed miRNAs of the 65 ranked by average log2FC values (Table 3).

**Table 3.** 21 miRNAs with significant differential expression and high patient frequencies (at least 60%) in tumor samples from each of eight or more cancers compared to non-tumorous samples

| miRNA<sup>a</sup> | Cancer type | Mean log2FC | Mean freq (strFC)<sup>b</sup> | Expression change |
| --- | --- | --- | --- | --- |
| **hsa-mir-1269a** | LUSC, LUAD, LIHC, UCEC, STAD, BRCA, HNSC, PRAD | 5.18 | 64.58% | Up |
| **hsa-mir-210** | LUSC, LUAD, BLCA, KIRC, UCEC, BRCA, HNSC, KIRP | 3.68 | 48.13% | Up |
| hsa-mir-301b | UCEC, LUSC, BLCA, LUAD, BRCA, ESCA, HNSC, KIRC, STAD, KIRP | 2.71 | 56.53% | Up |
| hsa-mir-183 | UCEC, BLCA, LIHC, KICH, BRCA, LUSC, LUAD, STAD, KIRP, PRAD, HNSC, THCA | 2.68 | 48.13% | Up |
| hsa-mir-3648 | LUSC, UCEC, ESCA, LUAD, BLCA, HNSC, PRAD, STAD, BRCA | 2.57 | 52.53% | Up |
| hsa-mir-3687 | BLCA, LUSC, LUAD, STAD, UCEC, HNSC, BRCA, PRAD | 2.56 | 52.91% | Up |
| hsa-mir-96 | UCEC, BLCA, KICH, BRCA, LIHC, LUSC, LUAD, STAD, PRAD, KIRP, THCA, HNSC | 2.56 | 49.69% | Up |
| hsa-mir-760 | LUSC, LIHC, LUAD, KIRC, KIRP, BRCA, UCEC, KICH, HNSC | 2.02 | 37.31% | Up |
| hsa-mir-4746 | UCEC, LIHC, ESCA, LUSC, KIRP, BRCA, HNSC, BLCA, KIRC, STAD, LUAD | 1.96 | 35.94% | Up |
| **hsa-mir-206** | BRCA, THCA, LUAD, KIRP, KIRC, LUSC, UCEC, KICH | -4.18 | 75.38% | Down |
| **hsa-mir-204** | KIRP, PRAD, LUAD, KIRC, STAD, THCA, HNSC, BLCA, BRCA, ESCA, KICH | -2.78 | 56.47% | Down |
| **hsa-mir-1-2** | KIRC, BRCA, KIRP, KICH, HNSC, ESCA, LUAD, LUSC, STAD, BLCA, UCEC | -2.74 | 59.78% | Down |
| **hsa-mir-133a-2** | PRAD, KIRC, LUAD, LUSC, KIRP, KICH, STAD, BLCA, UCEC | -2.72 | 59.94% | Down |
| **hsa-mir-133b** | THCA, PRAD, HNSC, KIRC, LUAD, KIRP, LUSC, KICH, STAD, BLCA, UCEC | -2.65 | 57.66% | Down |
| hsa-mir-133a-1 | THCA, KIRC, BRCA, HNSC, KIRP, ESCA, LUAD, KICH, LUSC, STAD, BLCA, UCEC | -2.6 | 59.94% | Down |
| hsa-mir-1247 | KIRC, LUSC, HNSC, LIHC, LUAD, THCA, KIRP, BRCA, KICH, BLCA, UCEC | -2.38 | 56.31% | Down |
| hsa-mir-139 | KIRC, THCA, LUAD, LIHC, HNSC, STAD, KIRP, LUSC, BRCA, BLCA, UCEC | -2.21 | 55.01% | Down |
| hsa-mir-145 | HNSC, THCA, LUAD, LIHC, LUSC, KICH, BRCA, STAD, KIRP, BLCA, UCEC | -1.73 | 39.98% | Down |
| hsa-mir-195 | LIHC, THCA, STAD, KICH, BRCA, HNSC, LUSC, LUAD, KIRP, BLCA, UCEC | -1.7 | 39.91% | Down |
| hsa-mir-99a | THCA, KICH, LIHC, LUAD, LUSC, BRCA, HNSC, UCEC | -1.63 | 33.44% | Down |
| hsa-let-7c | THCA, LIHC, KICH, LUSC, LUAD, HNSC, BRCA, UCEC | -1.55 | 34.02% | Down |

Molecular mechanisms for the newly identified hsa-mir-1269a have rarely been published for the cancer types we report here, but Min et al. have recently found a single nucleotide variation in hsa-mir-1269a that could contribute to the occurrence of hepatocellular carcinoma[38]. In our results, hsa-mir-1269a has log2FC value consistently larger than 2.20 in each of its eight associated cancer types (average log2FC = 5.18), with the average frequency of affected patients, greater than 64.58% for a strict fold change requirement (FC > 5 for each patient). Other members of the 21 key SDEmiRNAs (Table 3) include hsa-mir-183 and hsa-mir-96, which belong to the mir-183-96-182 cluster, which functions to induce cell proliferation and cancer development[39]. Notably, unlike the other two miRNAs from the same cluster, hsa-mir-182 only shows significantly differential expression but not high patient frequencies. Considering our relatively stringent criteria on FC values (FC > 2 or FC < 0.5), the changes of hsa-mir-182 in most patients are not as strong as that of hsa-mir-183 and hsa-mir-96.

Overall, our study groups 14 cancer types with potential different molecular mechanisms of development based on the distribution of SDEmiRNAs, as well as proposes some SDEmiRNAs previously indicated to play a role in distinct cancer types may function similarly in the development of additional cancer types, and the classification of small miRNA families into larger ones. Moreover, the results correspond well with those of previously published studies, and include evaluation of the 21 key SDEmiRNAs with high patient frequencies in a comprehensive pan-cancer analysis. Newly identified miRNAs hsa-mir-4746, hsa-mir-3648, hsa-mir-3687, and hsa-mir-1269a could also be important factors in multiple cancer types, although their relations to cancer were rarely reported by previous studies due to their low expression levels compared to others.

### SDEmiRNA targets enrichment and functional analysis

Each of the 65 SDEmiRNAs discussed above (with the same significantly differential expression trend in eight or more cancer types) participates in some form of molecular regulation, either inducing or inhibiting cell proliferation and cancer metastasis in different cancer types. In order to better understand the general pathways or biological processes by which SDEmiRNAs contribute to the occurrence and development of multiple cancer types, we extracted experimentally validated non-tissue-specific targets for those SDEmiRNAs without tissue or cancer information, as well as tissue-specific targets.

#### Distribution and functional analysis of non-tissue-specific targets of 65 SDEmiRNAs

60 of 65 selected miRNAs were found with a total 9,896 targets and a total of 29,087 miRNA-target interactions (the other five of which had no available target information). Among the target genes, 901 genes are considered to be cancer census genes from COSMIC or cancer biomarkers from EDRN, corresponding to 58 miRNAs. Cancer census genes are those genes with substantial evidence for their relations to cancer from published research[40]. hsa-mir-21 alone regulates 471 cancer-related genes, and hsa-mir-93 regulates 234 cancer-related genes. We then identified a subset of 27 cancer-related genes that are targets of more than 10 miRNAs, and among all miRNAs involved in regulating these 27 genes, 36 are found that target four or more. As shown in Figure 3, some of the cancer-related genes such as CCND2 (G1/S-specific cyclin-D2) (oncogene), MDM2 (E3 ubiquitin-protein ligase Mdm2) (oncogene), CDKN1A (Cyclin-dependent kinase inhibitor 1) (tumor suppressor), and SMAD4 (Mothers against decapentaplegic homolog 4) (tumor suppressor) could be regulated by multiple miRNAs, and the numbers of targets of hsa-mir-106b, hsa-mir-19a, hsa-mir-195, hsa-mir-130b, and hsa-mir-21 are greater than that of others.

**Figure 3.**
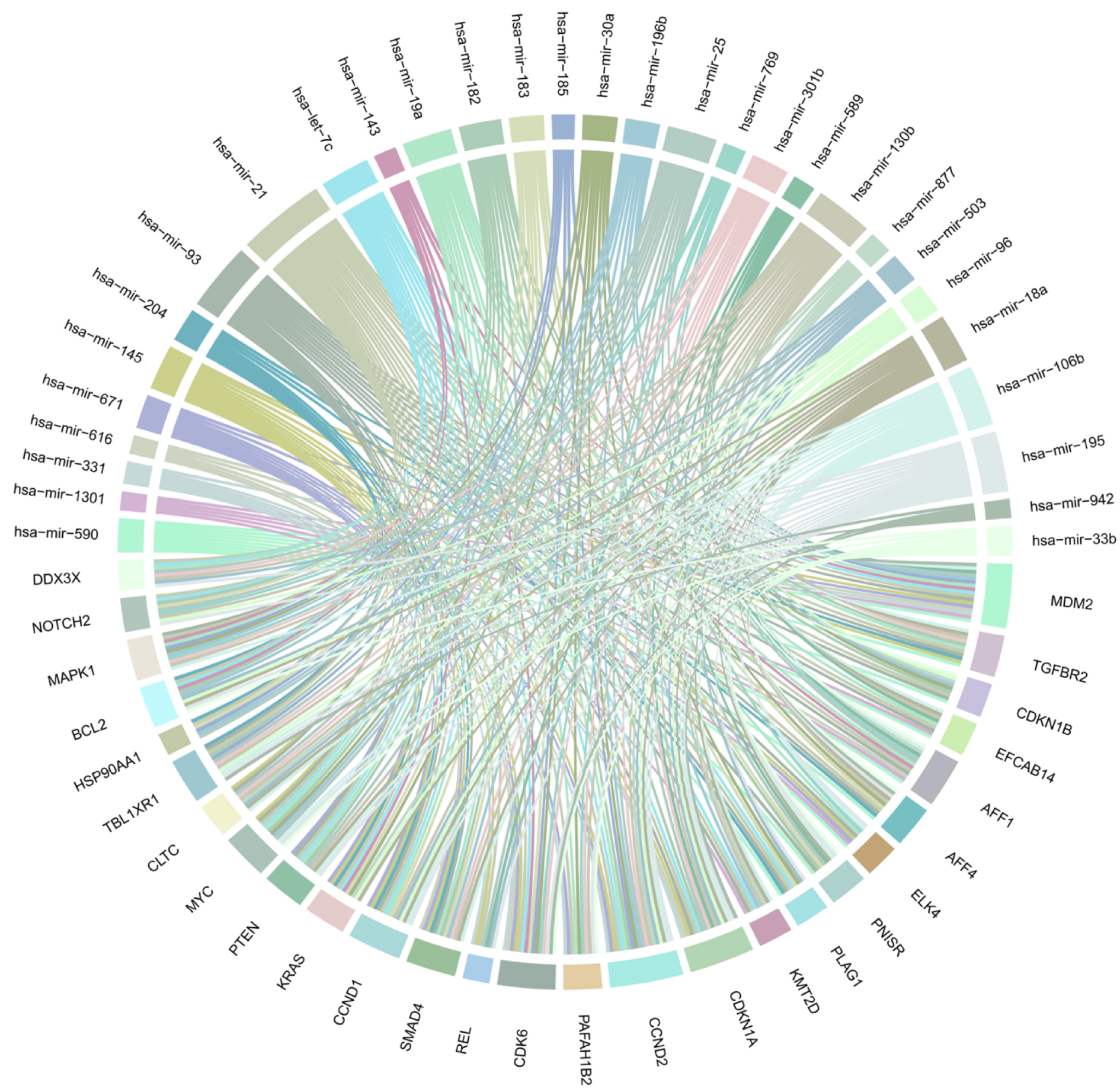
Non-tissue-specific cancer-related genes which are targets of more than 10 miRNA types. Each unique color represents one miRNA with at least four targets and the ribbon width suggests the interactions between the miRNAs and target genes. 27 cancer-related genes are targets of more than 10 miRNAs, and among all miRNAs involved in regulating these 27 genes, 36 target four or more cancer-related genes. Some of the cancer-related genes could be regulated by multiple miRNAs, and the numbers of targets of some miRNAs are greater than those of others. These miRNAs play vital roles in cancer progression via regulating cancer-related genes.

For the 21 key SDEmiRNAs with high patient frequencies (FC > 2 or < 0.5) (Table 3), as well as hsa-mir-4652, which has the largest log2FC values and more than 60% of patient frequency in seven cancers, 18 were found to have 428 cancer-related ones out of their 3,987 non-tissue-specific targets. Fifteen of the 21 key SDEmiRNAs regulate 25 cancer-related genes. Eleven of these SDEmiRNAs function in enzyme binding (p value = 8.30e-03, by STRING), lending credence to their regulatory roles in cancer development. Well-studied miRNAs such as hsa-mir-195, hsa-let-7c, and hsa-mir-183 have more than 70 cancer-related targets each. In particular, hsa-mir-1269a has 57 (five cancer-related), hsa-mir-4652 has 133 (12 cancer-related), hsa-mir-4746 has 54 (six cancer-related), hsa-mir-3648 has 14 (one cancer-related), and hsa-mir-3687 has 14 (no cancer-related) non-tissue-specific targets. Our analysis by IPA suggests these targets participate in various important networks and cellular processes (Table 4). For cancer-related targets of hsa-mir-1269a and hsa-mir-4652, most of them are involved in cell survival, such as cell cycle progression, cell migration, and apoptosis (Figure 4A-B), while targets of hsa-mir-4746 are less condensed and function mostly in more specific processes, such as cell death of cervical cancer cell lines (Figure 4C). The one cancer-related target of hsa-mir-3648 is CCND1 (G1/S-specific cyclin-D1) involved in cell cycle control.

**Table 4.** Newly identified SDEmiRNAs and the potential networks and pathways involved in cancer

| **SDEmiRNA** | **Number of non-tissue-specific target** | **Networks and Pathways** |
| --- | --- | --- |
| hsa-mir-1269a | 57 | Molecular transport, RNA trafficking, cell death and survival<br>Lipid metabolism, nucleic acid metabolism<br>Cellular assembly and organization, DNA replication, recombination, and repair, cell morphology<br>Cardiovascular system development and function, organismal development |
| hsa-mir-4652 | 133 | Cellular development, cellular growth and proliferation<br>Organ Morphology, organismal development, organismal injury and abnormalities<br>Cardiovascular system development and function, skeletal and muscular system development and function<br>Immune response |
| hsa-mir-4746 | 54 | Skeletal and muscular disorders, organismal development<br>Cellular assembly and organization, nervous system development and function, tissue morphology<br>Cellular development, cellular growth and proliferation<br>Intracellular membrane-bounded organelle |
| hsa-mir-3648 | 14 | Connective tissue disorders, development disorder, endocrine system disorders |
| hsa-mir-3687 | 14 | Cellular development, cellular growth and proliferation, cell cycle |

**Figure 4.**
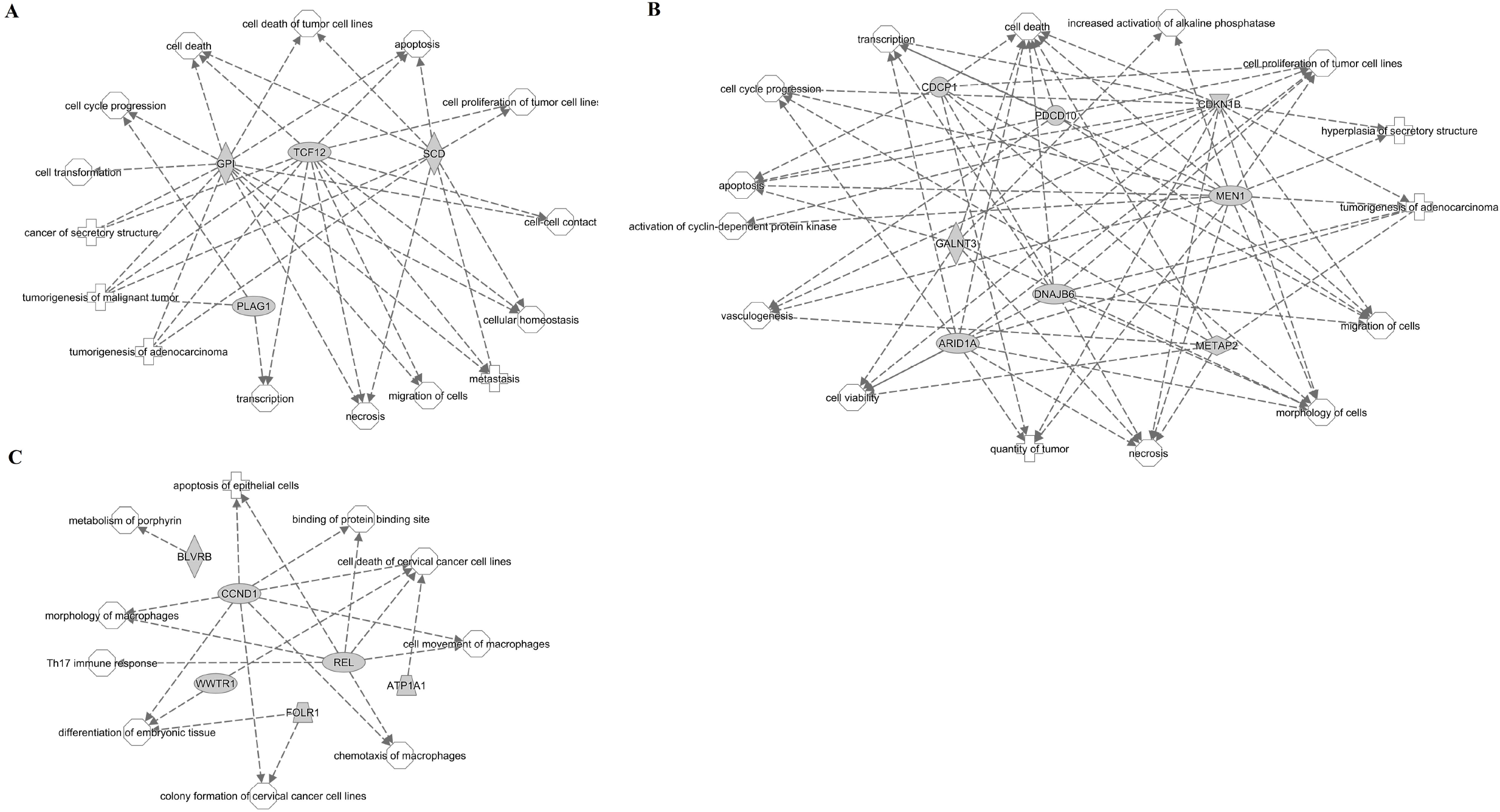
Networks and pathways of non-tissue-specific cancer-related targets of hsa-mir-1269a, hsa-mir-4652, and hsa-mir-4746. **A.** Four out of five non-tissue-specific cancer-related targets of hsa-mir-1269a with grey shapes participate in 15 cellular processes with white shapes. Each arrow represents one correlation between a target and a process. GPI, TCF12, and SCD are involved in most of the processes, compared to PLAG1. **B.** Network of eight out of 12 cancer-related targets of hsa-mir-4652. These targets all participate in multiple cellular processes. **C.** Network of all six cancer-related targets of hsa-mir-4746. Except BLVRB, other targets share similar processes with each other. Based on the important roles of these cellular processes in cancer development, the three newly identified SDEmiRNAs may function in cancers through these processes.

By targeting a wide range of cancer-related genes, those 65 SDEmiRNAs, especially 21 key SDEmiRNAs, participate in multiple processes, contributing to cancer development. Moreover, those newly identified SDEmiRNAs could be equally important with others, despite their relatively lower expression levels. They may be able to achieve their contributions to cancer through regulating targets within small regions or participating in short-term stress response, which requires less dosage.

#### Distribution of tissue-specific targets of 65 SDEmiRNAs and functional analysis of them

We also collected tissue-specific targets for the 65 SDEmiRNAs (significantly expressed in at least eight cancer types) (Table S3) and explored the potential molecular mechanisms of those tissue-specific targets in light of their observed expression trends and biological processes in which they participate. All selected miRNAs and their targets were mapped to 11 cancer types by DOIDs. A total of 9,572 target genes were found, of which 7,325 were mapped to reviewed UniProtKB accession numbers (Table S4). 7,186 of the implicated significant, reviewed target genes are included in BioXpress v3.0, which reports genes with significant expression changes derived from paired tumor and non-tumorous data from TCGA: currently, there are 5,971 genes showing significant differential expression in multiple cancer types in BioXpress (Table S5). miRNA target genes for lung cancer and kidney cancer are a combination of those identified for TCGA cancer types LUAD and LUSC, KICH and KIRP and KIRC, respectively, and have the same expression trend across these TCGA cancer types.

Mapping to Gene Ontology (GO) terms and PANTHER pathway IDs resulted in retrieval of 12,467 GO terms and 130 PANTHER terms. According to the screening conditions described in the methods section, 99 records were selected in Table S6 (22 from PANTHER terms and 77 from GO terms). The criteria of the screening were such that we expected to lose potentially valuable GO or PANTHER terms, but we chose to follow a conservative method to reduce possible false positive results. The most common molecular functions these miRNAs and their targets participate in cancers are negative regulation of ERK1 and ERK2 cascade (GO:0070373), canonical Wnt signaling pathway (GO:0060070), and positive regulation of sequence-specific DNA binding transcription factor activity (GO:0051091). Each of these functions are involved in ten types of cancer, with more than 40 SDEmiRNAs, and more than 30 targets, respectively. This result not only is consistent with Gosline et al.[41], which suggested the strong regulation of miRNAs on transcription factors in mice, but also suggests most of these SDEmiRNAs could promote the activity of transcription factors.

These terms suggest the participation in regulation of cell proliferation and apoptosis through different mechanisms, and frequently implicated cancer pathways were found in our list, including the p53 pathway, Wnt signaling pathway, TGF-β/SMAD signaling pathway, and others. We also found that the occurrence and development of these cancer types may share similar molecular mechanisms with other diseases or processes, including Alzheimer’s disease (AD) (with involved amyloid secretase and presenilin pathways), for which the product the key gene implicated in AD, the amyloid precursor protein, is over-expressed in multiple cancer types[42] and other neurodegenerative diseases.

For the 21 key SDEmiRNAs (with both significant differential expression and at least 60% patient frequencies in at least eight cancer types) (Table 3), as well as hsa-mir-4652, 19 were found to have tissue-specific targets. We identified the top one or two genes that are targets of the most SDEmiRNAs in each cancer type. In breast cancer, CCND1 and IGF1R (Insulin-like growth factor 1 receptor) are found to be the targets of nine and seven key SDEmiRNAs. IGF1R is also the most popular target of eight SDEmiRNAs in lung cancer, and of seven SDEmiRNAs in head and neck cancer. CCND1 promotes G1/S transition during mitotic cell cycle by regulating CDK4 (cyclin-dependent kinase 4), and is up-regulated in various cancer types including breast cancer[43, 44]. IGF1R is also a well-studied oncogene that allows cell proliferation and growth, and inhibits apoptosis[45]. Our RNA-seq analysis in BioXpress v3.0 shows that CCND1 is significantly over-expressed in BRCA, while IGF1R has significant overexpression in BRCA, LUSC, and HNSC. The roles of CCND1 and IGF1R could be vital and essential, compared to other cancer-related ones, for the development of these cancer types, considering such huge compensation mechanisms over their regulation.

In addition, in breast cancer hsa-mir-1269a has one target (EN2, Homeobox protein engrailed-2) which is a sequence-specific DNA binding protein positively regulating transcription from RNA polymerase II promoter. hsa-mir-4652 has two targets (GALNT3, EIF4EBP1) in lung cancer. GALNT3 (Polypeptide N-acetylgalactosaminyltransferase 3) functions as an iron binding enzyme in glycosylation, while EIF4EBP1 (Eukaryptic translation initiation factor 4E-binding protein 1) binds to IRES-dependent translational initiation to repress protein translation and participates in insulin receptor signaling pathway and G1/S transition of mitotic cell cycle (Table 4). hsa-mir-4652 has another target (H2AFX, Histone H2AX) involved in DNA and histone binding in head and neck cancer. CancerMiner predicted 12 targets of hsa-mir-3648 in head and neck cancer (Table S4). Their functions include ATP binding, cell cycle control, lipid metabolism, protein folding, and nucleobase biosynthesis. hsa-mir-4746 has a common target, PPM1D (Protein phosphatase 1D), in both breast and lung cancer. Because of its function in protein dephosphorylation, it is involved in G2/M transition of mitotic cell cycle and suppression of cell proliferation in cancer[46]. The importance of the functions of their target genes in gene/cellular regulation suggests abnormally expressed hsa-mir-1269a, hsa-mir-4652, and hsa-mir-4746 may be able to contribute to the development of corresponding cancer types.

### Construction of regulatory networks of SDEmiRNAs and their targets

Suppression of a negative regulator will result in up-regulation of the regulator’s target molecule. Similarly, because miRNAs function to suppress gene expression, miRNA inhibition of a negative regulator will also result in up-regulation of the target molecule. There are some cases in which a miRNA can have two targets such that one target regulates the other. In these cases, up-regulated targets of over-expressed miRNAs or down-regulated targets of under-expressed miRNAs are considered to be indirect targets: miRNAs either inhibit the negative regulators or activate the positive regulators of the differentially expressed target genes, respectively.

Across eight cancer types, there are 34 SDEmiRNAs that regulate 14 potential target genes in the angiogenic patterning of blood vessels (GO:0001569), which is essential for establishing abnormal tumor blood vessels[47]. Our results here show hsa-mir106b, hsa-mir-93, and hsa-mir-21, which are over-expressed in six cancers and target eight genes (out of the 14 possible targets), play important roles in tumor blood vessel generation. All three SDEmiRNAs directly target one key tumor suppressor gene, TGFBR2 (TGF-beta receptor II)[48], leading to its underexpression. Unlike over-expressed miRNAs with a relatively large number of targets, most under-expressed miRNAs only target central genes of angiogenesis, including TGFBR2 (indirect), VEGFA (vascular endothelial growth factor A) (direct), and CTNNB1 (beta-catenin) (direct). These trends could suggest that during patterning of blood vessels in these eight cancer types, most tumor suppressive miRNAs regulating angiogenic central genes are repressed, while the three oncogenic miRNAs above activate a variety of relevant genes, efficiently promoting the tumorigenic process.

To further elucidate the underlying mechanisms of miRNA involvement in cancer, we can construct a network of miRNA regulation of signaling pathways across different cancer types. The PI3-kinase (PI3K) pathway (PANTHER: P00048) is one of the most important pathways that participates in the regulation of cancer development and growth[49]. PTEN, an important tumor suppressor, is a negative regulator of the PI3K pathway[50]. As shown in Figure 5, eight target genes of 16 miRNAs are involved in the PI3K pathway across seven cancer types, based on KEGG (Release 79.1, http://www.genome.jp/kegg/) and studies from Waugh MG[51], Ying Z et al.[52], and Yun YR et al.[53]. miRNAs involved in regulating FGFR family members were also shown in Figure 5, adding another potentially important miRNA, hsa-mir-381, to the pathway.

**Figure 5.**
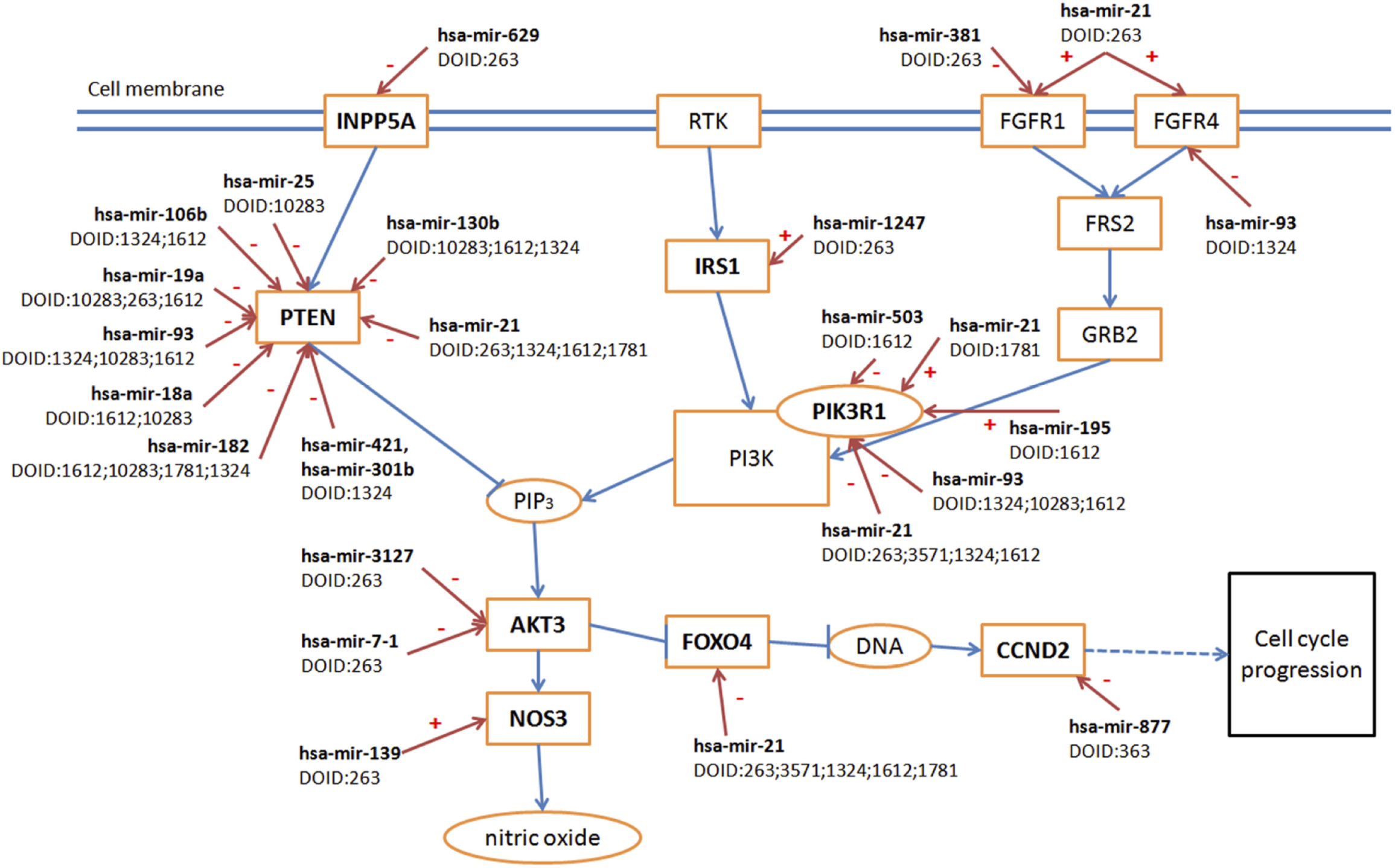
Regulatory network of miRNA on PI3-kinase pathway. Blue arrows indicate the regulation among genes in the PI3-kinase pathway (arrows with vertical short line segments represent inhibition role of up-stream genes). Red arrows indicate regulation interaction between miRNAs and PI3-kinase pathway. Minus sign suggests inhibition, and plus sign promotion. Eight target genes of 16 miRNAs are involved in the PI3K pathway and regulation of PTEN gene across seven cancer types. Different miRNAs in different cancers could regulate the same molecular processes through targeting their components. These miRNAs still could function similarly across at least eight cancer types.

Most over-expressed miRNAs silence PTEN across different cancer types, increasing the signal passing from PIK3 to AKT3, and some miRNAs regulate PI3KR1 (phosphoinositide-3-kinase regulatory subunit 1, a gene of PI3K Class I). hsa-mir-21, for instance, is a well-known, over-expressed miRNA in various cancer types. Our results in Figure 5 indicate that hsa-mir-21 could inhibit PI3KR1 in breast, liver, kidney, and lung cancer, while it promotes PI3KR1 in thyroid cancer, suggesting that for cancer types other than thyroid cancer, other members of the PI3K classes of enzymes, may play more important roles in cancer progression. It suggests different SDEmiRNAs could still regulate the same molecular processes in different cancer types, even if their target genes may vary from each other. This network integrates current discoveries on SDEmiRNAs, their targets, and related functions, displays SDEmiRNA regulation in multiple cancer types, and therefore, provides a straightforward view of what can be done in future analysis. Similar networks could be constructed based on other results reported herein, and could ultimately lay the foundation for further studies regarding the role of miRNA expression in cancer.

## Materials and Methods

### Data

#### miRNA sequencing data

This study started with miRNA sequencing data integration and filtering, and performed expression analysis and downstream functional analysis (Figure 6). All miRNAseq data (both counts of raw reads and Reads Per Million miRNA mapped (RPM) values, annotated to miRNA names level) were downloaded from TCGA (Release-2016-03) (http://cancergenome.nih.gov) by using TCGA-Assembler[54] with default parameters (Figure 6). Data was then divided into two groups: paired data—data with both tumor and adjacent non-tumorous samples from the same patient, and tumor-only expression data—data coming exclusively from all tumor samples in all TCGA cancer types with miRNAseq data. Paired data were limited to those generated on the HiSeq platform which generated enough matched samples than other platforms, whereas tumor-only expression data were focused on data generated either on HiSeq or Genome Analyzer (GA) of Illumina[55] systems, analyzed in separate groups for data from each platform. Two additional projects (MALY, Malignant Lymphoma done by Germany, and OV, Ovarian Cancer done by Australia, Release 20), available only through the International Cancer Genome Consortium (ICGC, https://icgc.org/) and generated by Illumina-HiSeq system, were imported into the tumor-only expression dataset. Fold change value (FC) was calculated by (RPM)^tumor^ / (RPM)^non-tumor^ for one miRNA in one cancer type. Each miRNA was mapped to its corresponding HGNC ID (HUGO Gene Nomenclature Committee, Issued in Autumn/Winter 2015: http://www.genenames.org), miRBase ID (Release 21)[1], and Ensembl ID (Release 82)[56] using customized python (v2.7) and R (3.3) scripts.

#### Databases for Validation of miRNA

To validate the selected differentially expressed miRNAs, we used four experimentally validated databases generated from literature mining: HMDD (v2.0)[57], Mir2disease (2008)[58], miRCancer (March2016)[59], and miRiaD (2016)[60]. We also applied our literature mining tool, DEXTER, to extract miRNA-cancer correlations from publications[61]. The basic search term is “[miRNA name] is [over-expressed / under-expressed / significantly higher / lower] in [cancer tissue/sample] compared to [normal / non-tumorous sample]”. This tool is currently being benchmarked, and the results were manually curated before reported as validate records. The cancer terms for each reported interaction between miRNA and specific cancer types were then mapped to Disease Ontology[62] cancer slim in order to unify the integrated dataset[63]. We also incorporated information regarding miRNA expression changes in cancer types (up/down-regulation) from these databases, when available.

#### miRNA Targets

Five experimentally validated miRNA target databases generated from literature mining were used in our study: miRTarbase (Release 6.0)[64], miRecords (Apr 2013)[65], miRWalk 2.0 (miRNA-target interaction databases)[66], miRTex (May 2015)[67], and Tarbase-DIANA lab v6.0[68]. These databases were generated by different algorithms but with a common motive, which is to extract miRNA-target-cancer/tissue interactions from published papers. Most of these published papers were validating the interactions or stating that the interactions were commonly accepted by previous studies. CancerMiner (1.0), a database for tissue-specific miRNA target predictions based on TCGA data[69], was also included in the integrated miRNA target database for this study. This database was built upon multivariate linear model for expression association of miRNA and mRNA from TCGA, and rank-based association recurrence score that showed the possible association of miRNA-target interactions and cancer types [69].

GO (Gene Ontology) IDs[70] and PANTHER pathway IDs and terms were downloaded from UniProt (http://www.uniprot.org/, Release 2016_08) and PANTHER ((Protein ANalysis THrough Evolutionary Relationships, Release 11.0: http://www.pantherdb.org/)[71] for each target in the resulting database.

### Data Normalization and Differential Expression Analysis

We used DESeq2 version 1.8.1 [26] and edgeR version 3.10.2[24] (R packages) to normalize paired miRNA sequencing data and to analyze the differential expression of miRNAs in each cancer type. These tools are both based on a negative binomial distribution assumption[27, 72], and both tools have been proven to generate normalized data of high quality and to perform similarly in differential expression analyses, as well as in fold change estimates[73, 74]_ENREF_19_ENR EF_23. DESeq2 fits a generalized linear model for each miRNA with the fold change estimate shrunken by empirical Bayes[26]. A Wald test was used in DESeq2 for statistical analysis and calculating *p* values for the significance of differentially expressed miRNAs in each cancer type. edgeR, however, uses the TMM method for normalization of sequencing data. To estimate parameters for the negative binomial model, we applied common dispersion, trended dispersion, and tagwise dispersion for our multi-factor cases. These dispersions were shrunken by an empirical Bayes procedure towards a consensus value[24], and edgeR applied Fisher’s Exact Test to determine differential expression[24]. False discovery rate (FDR) was reported by edgeR for the significance of multiple comparisons conducted. The input data for the two tools were raw read counts (without normalization), excluding any miRNAs with no expression among all patients. For both tools, we designed the model matrix with two categories, one with levels “cancer” and “non-tumor”, the other with TCGA patient IDs. The cutoff for the significant adjusted p-value of each gene is 0.05/n (using Bonferroni’s Approach, where n is the total number of expressed miRNAs in each cancer type), considering the difference of cancers. miRNAs with significant adjusted p-values generated by both DESeq2 and edgeR were considered to be differentially expressed in a given cancer type. Results from differential expression analysis for paired and tumor-only expression data were integrated into BioXpress version 3.0[75]. The Heatmap with clustering for significantly differentially expressed miRNAs across different cancer types was generated by R package *gplots* 3.0.1.

### Survival Analysis

miRpower[36] was used to suggest the influence of miRNAs on patient survival rate in breast cancer. This platform is continuously updated with TCGA datasets, and has been cited by dozens of studies. By providing “auto select best cutoff”, miRpower is able to optimize the cutoff of the patient groups for different investigated miRNAs. In our study here, TCGA was selected as the only dataset to be considered, and therefore, miRNAs from breast cancer (BRCA) were under survival analysis (1,061 TCGA patients in total).

### miRNA Target Data Integration and Enrichment

After extracting all targets for selected miRNAs (those significantly over- or under-expressed in at least eight cancer types), we separated these targets into two categories: tissue-specific targets (records clearly including the miRNA-target connection in a specific tissue or cancer type), and non-tissue-specific targets (records containing only a miRNA-target connection). Tarbase provided the majority of records for the tissue-specific target group. We excluded associations derived from microarray studies and integrated those records with cancer metastasis annotation. Reported regulatory changes (up/down) of miRNAs were also retrieved. Selective records from miRWalk 2.0 were included in the tissue-specific target group if the records were involved in miRNAs-organ interactions or miRNA-gene-OMIM interactions generated by miRWalk text-mining search on PubMed database. To include more tissue-specific targets, we also integrated data from a predicted database, CancerMiner, which applied mRNA and miRNA expression, DNA copy number, and promoter methylation data from TCGA to predict miRNA-mRNA-cancer type associations[69]. CancerMiner data were included only for records not found in Tarbase or miRWalk. Expression changes of all tissue-specific target genes in corresponding cancer types were extracted from BioXpress v3.0[75]. For the non-tissue-specific target group, we first excluded those targets that were also in the tissue-specific target section. Some records from Tarbase that do not contain tissue or cancer information were included. Gene names of the records from miRTex were first manually curated and mapped to synonyms available in UniProtKB, and records with miRNA-gene associations were selected while miRNA-miRNA interactions were excluded. miRNA-gene targets from miRWalk 2.0 and records from miRTarbase and miRecords were all classified into the non-tissue-specific target group. All cancer types from those databases were mapped to DOIDs.

Tissue-specific target genes were mapped to UniProtKB accession numbers[76], maintaining only those genes that have been manually annotated and reviewed in UniProtKB/Swiss-Prot. For each GO or PANTHER term, we merged all the target genes involved and their regulating miRNAs. In order to reduce false positive errors and to exclude highly generic terms of GO, GO terms of genes associated more than eight cancer types and fewer than 50 target genes were selected. We also collected non-tissue-specific targets that are considered to be cancer census genes from COSMIC (version 78)[77] and biomarkers from EDRN (Downloaded in Feb, 2016, https://edrn.nci.nih.gov/). Enrichment analysis of these nontissue-specific targets was done using STRING (version 10.0)[78] and Ingenuity Pathway Analysis (IPA) (March 2017).

## Conclusion

Different cancer types have unique expression patterns of significantly differentially expressed miRNAs (SDEmiRNAs). Our study here sought to identify prevalent expression-based or mechanistic features of SDEmiRNAs associated with cancer development, even though the analysis here did not separate cancer samples based on clinical information, such as tumor stage or patient metadata. Applying our miRNA analysis findings, we propose to narrow down the number of miRNAs needed to classify the 14 cancer types, as well as cell type cancers from the same location. Moreover, based on the inhibition role of miRNA, development of different cancer types may rely on either more tumor suppressors or more oncogenes, resulting in three categories of cancers. We also identified 81 unique SDEmiRNAs with significantly differential expression in only one out of the 14 cancer types: these SDEmiRNAs might have unique functions in each cancer, even including effect on patient survival rates, although most of them have not been studied in their corresponding cancer type and their expression levels are lower than well-studied ones.

We then identified a set of 21 key SDEmiRNAs (including four relatively newly identified ones, hsa-mir-4746, hsa-mir-3648, hsa-mir-3687, and hsa-mir-1269a) that have high enrichment in patients of at least eight cancer types, as well as hsa-mir-4652 with high enrichment in seven cancer types. These key SDEmiRNAs may have much wider and more similar regulatory mechanisms across multiple cancer types than what researchers have found in one or two cancers so far. Our study also shows the necessary to study further on those newly identified ones, which might be equally important with others. To explore the possible molecular mechanisms of these key SDEmiRNAs, enrichment analysis of the SDEmiRNAs and their targets in different cancer types was applied, which suggested that different miRNAs participate in a limited amount of cancer-associated molecular processes/pathways through regulating different targets. This means they may have similar mechanisms in development of these cancer types.

The construction of regulatory network of SDEmiRNAs with their experimentally validated targets combines results of current studies and shows the potential research directions for further analysis. Our study here provides a rationale for the continued exploration and validation of the functional roles of key SDEmiRNAs in cancers, and therefore, pushes all the results on BioXpress to allow further analysis.

## Authors’ Contributions

R.M. supervised the study. Y.H. and R.M. contributed to conception and design. Y.H., Q.W. and C.Y. were involved in methodology and acquisition of data. Y.H. analyzed and interpreted data. Y.H., R.K., S.G. and V.S. participated in constructing database or data support. Y.H., H.D., S.G., R.K., V.S., Q.W., C.Y. and R.M. wrote and reviewed the manuscript.

## Acknowledgments

The authors thank Yu Fan and Ting-Chia Chang for helpful discussion. Yu Hu earned an MS in the Bioinformatics Track of the Bioinformatics and Molecular Biochemistry Program at the George Washington University. This work is from a dissertation to be presented to the above program in partial fulfillment of the requirements for the Master of Science degree.

## Additional Information

Results of miRNA differential expression analysis are available in: https://hive.biochemistry.gwu.edu/bioxpress

---

**Figure 6. Cancer-related significantly differential expression of miRNA workflow**

Paired miRNAs (tumorous and adjacent non-tumorous samples) were retrieved from TCGA database and analyzed by DESeq2 and edgeR for differential expression. We then screened out those significantly differentially expressed miRNAs (SDEmiRNAs). After validation by literature mining databases and our text-mining tool, DEXTER, we proposed the possibility to use these SDEmiRNAs in classification of cancer types or cancer subtypes, and screened out unique SDEmiRNAs for one single cancer. We also kept those key SDEmiRNAs significantly expressed in at least eight cancers with high patient frequencies. Moreover, we collected both tissue-specific targets and non-tissue-specific targets from experimentally validated databases, and did functional/enrichment analysis with those key SDEmiRNAs.

## Supplementary Materials

**Figure S1 Webpage of each miRNA in BioXpress v3.0**

Results of differential expression analysis of human whole miRNAs are shown by both table and figures in BioXpress v3.0. Each miRNA is also mapped to corresponding Ensembl, miRbase, RefSeq, and HGNC identifiers to allow users to search. Cancer types are also mapped to Disease Oncology IDs.

**Figure S2 Identification of different cell types within one organ cancer by miRNA**

**A.** Lung cancer - there are 190 significantly differentially expressed miRNAs with Δlog2FC > 1 between LUAD (Lung adenocarcinoma) and LUSC (Lung squamous cell carcinoma), including 50 having the same expression change in these two cell types. The remaining 140 miRNAs with opposite expression patterns are considered to be key factors underlying the differences between LUAD and LUSC. **B-D.** Kidney cancer - Pairwise comparisons were made between KIRP (Kidney renal papillary cell carcinoma) and KIRC (Kidney renal clear cell carcinoma) (**B**), KICH (Kidney Chromophobe) and KIRP (**C**), and KIRC and KICH (**D**). There are 330 significantly differentially expressed miRNAs with Δlog2FC > 1 between any two of the three subtypes. miRNA types and order are preserved across the three comparisons of kidney cancer. From the comparison, the development of KIRC and KIRP may be mechanistically closer with each other, compared to the development of KICH.

**Figure S3 Significantly differentially expressed miRNAs that are over-expressed in at least 80 percent of patient or under-expressed per cancer**

Log2FC values were used to distinguish the over- or under-expression of miRNAs per patient. Green bars with values larger than 80 percent indicate miRNAs are over-expressed in at least 80% of patient, while red bars with values larger than 80 present miRNAs (green bars lower than 20) that are under-expressed in at least 80% of patient per cancer. In the figure, some cancers have more over-expressed miRNAs than under-expressed ones, while others have the opposite trend. This suggests different cancer types can have different enrichment patterns of differentially expressed miRNAs, which implies the potential existence of different mechanisms in their occurrence and development.

**Figure S4 Survival analysis of three SDEmiRNAs specific to BRCA, hsa-mir-4784, hsa-mir-1262, and hsa-mir-320c-1 by miRpower**

**A.** Kaplan-meier plot of hsa-mir-4784. Patients with under-expressed hsa-mir-4784 show higher survival rate than the ones with over-expressed hsa-mir-4784. Consistence with our result suggests significant over-expression of hsa-mir-4784 may be vital for BRCA development. **B.** Kaplan-meier plot of hsa-mir-1262. **C.** Kaplan-meier plot of hsa-mir-320c-1. Patients with underexpressed hsa-mir-1262 and hsa-mir-320c-1 have lower survival rate compared to those with over-expressed ones. This result also fits our study of the significant under-expression of hsa-mir-1262 and hsa-mir-320c-1.

**Table S1 Unique miRNAs with significantly differential expression in one single cancer type of the 14**

**Table S2 Validation of a subset of 24 out of 90 significantly differentially expressed miRNAs by text mining tool DEXTER**

**Table S3 Significantly differential expressed miRNAs that are all over-expressed or all under-expressed in more than eight cancer types**

**Table S4 Tissue-specific targets of each selected miRNA per cancer type**

Targets are represented by UniProt Accession Numbers.

**Table S5 Significantly differential expressed miRNA in at least eight cancer types and their target counts for each cancer type**

**Table S6. Selected GO and PANTHER terms involved in miRNAs and their tissue-specific targets.** Cancer types are represented by DOIDs (Disease Ontology ID).

## References

[1] Kozomara A, Griffiths-Jones S. miRBase: annotating high confidence microRNAs using deep sequencing data. Nucleic Acids Res 2014;42:D68–73.

[2] Zempleni J, Baier SR, Howard KM, Cui J. Gene regulation by dietary microRNAs. Can J Physiol Pharmacol 2015;93:1097–102.

[3] Calin GA, Dumitru CD, Shimizu M, Bichi R, Zupo S, Noch E, et al. Frequent deletions and down-regulation of micro- RNA genes miR15 and miR16 at 13q14 in chronic lymphocytic leukemia. Proc Natl Acad Sci U S A 2002;99:15524–9.

[4] Takamizawa J, Konishi H, Yanagisawa K, Tomida S, Osada H, Endoh H, et al. Reduced expression of the let-7 microRNAs in human lung cancers in association with shortened postoperative survival. Cancer Res 2004;64:3753–6.

[5] Chan JA, Krichevsky AM, Kosik KS. MicroRNA-21 is an antiapoptotic factor in human glioblastoma cells. Cancer Res 2005;65:6029–33.

[6] Ma L, Teruya-Feldstein J, Weinberg RA. Tumour invasion and metastasis initiated by microRNA-10b in breast cancer. Nature 2007;449:682–8.

[7] Gonzalez-Quintana V, Palma-Berre L, Campos-Parra AD, Lopez-Urrutia E, Peralta-Zaragoza O, Vazquez-Romo R, et al. MicroRNAs are involved in cervical cancer development, progression, clinical outcome and improvement treatment response (Review). Oncol Rep 2016;35:3–12.

[8] Tutar L, Tutar E, Ozgur A, Tutar Y. Therapeutic Targeting of microRNAs in Cancer: Future Perspectives. Drug Dev Res 2015;76:382–8.

[9] Wang W, Luo YP. MicroRNAs in breast cancer: oncogene and tumor suppressors with clinical potential. J Zhejiang Univ Sci B 2015;16:18–31.

[10] Hayes J, Peruzzi PP, Lawler S. MicroRNAs in cancer: biomarkers, functions and therapy. Trends Mol Med 2014;20:460–9.

[11] Thomas J, Ohtsuka M, Pichler M, Ling H. MicroRNAs: Clinical Relevance in Colorectal Cancer. Int J Mol Sci 2015;16:28063–76.

[12] Khan MA, Zubair H, Srivastava SK, Singh S, Singh AP. Insights into the Role of microRNAs in Pancreatic Cancer Pathogenesis: Potential for Diagnosis, Prognosis, and Therapy. Adv Exp Med Biol 2015;889:71–87.

[13] Lu J, Getz G, Miska EA, Alvarez-Saavedra E, Lamb J, Peck D, et al. MicroRNA expression profiles classify human cancers. Nature 2005;435:834–8.

[14] Ren Z, Wang W, Li J. Identifying molecular subtypes in human colon cancer using gene expression and DNA methylation microarray data. Int J Oncol 2016;48:690–702.

[15] Sisti JS, Collins LC, Beck AH, Tamimi RM, Rosner BA, Eliassen AH. Reproductive risk factors in relation to molecular subtypes of breast cancer: Results from the nurses’ health studies. Int J Cancer 2016;138:2346–56.

[16] Gasparini P, Cascione L, Landi L, Carasi S, Lovat F, Tibaldi C, et al. microRNA classifiers are powerful diagnostic/prognostic tools in ALK-, EGFR-, and KRAS-driven lung cancers. Proc Natl Acad Sci U S A 2015;112:14924–9.

[17] Monroig-Bosque Pdel C, Rivera CA, Calin GA. MicroRNAs in cancer therapeutics: “from the bench to the bedside”. Expert Opin Biol Ther 2015;15:1381–5.

[18] Sethi S, Ali S, Sarkar FH. MicroRNAs in personalized cancer therapy. Clin Genet 2014;86:68–73.

[19] Peng F, Zhang Y, Wang R, Zhou W, Zhao Z, Liang H, et al. Identification of differentially expressed miRNAs in individual breast cancer patient and application in personalized medicine. Oncogenesis 2016;5:e194.

[20] Metpally RP, Nasser S, Malenica I, Courtright A, Carlson E, Ghaffari L, et al. Comparison of Analysis Tools for miRNA High Throughput Sequencing Using Nerve Crush as a Model. Front Genet 2013;4:20.

[21] Humphreys DT, Suter CM. miRspring: a compact standalone research tool for analyzing miRNA-seq data. Nucleic Acids Res 2013;41:e147.

[22] Lopes-Ramos CM, Habr-Gama A, Quevedo Bde S, Felicio NM, Bettoni F, Koyama FC, et al. Overexpression of miR-21-5p as a predictive marker for complete tumor regression to neoadjuvant chemoradiotherapy in rectal cancer patients. BMC Med Genomics 2014;7:68.

[23] Cobos Jimenez V, Willemsen AM, Bradley EJ, Baas F, van Kampen AH, Kootstra NA. Next-generation sequencing of microRNAs in primary human polarized macrophages. Genom Data 2014;2:181–3.

[24] Robinson MD, McCarthy DJ, Smyth GK. edgeR: a Bioconductor package for differential expression analysis of digital gene expression data. Bioinformatics 2010;26:139–40.

[25] Anders S, Huber W. Differential expression analysis for sequence count data. Genome Biol 2010;11:R106.

[26] Love MI, Huber W, Anders S. Moderated estimation of fold change and dispersion for RNA-seq data with DESeq2. Genome Biol 2014;15:550.

[27] Yang S, Guo L, Shao F, Zhao Y, Chen F. A Systematic Evaluation of Feature Selection and Classification Algorithms Using Simulated and Real miRNA Sequencing Data. Comput Math Methods Med 2015;2015:178572.

[28] Funamizu N, Lacy CR, Parpart ST, Takai A, Hiyoshi Y, Yanaga K. MicroRNA-301b promotes cell invasiveness through targeting TP63 in pancreatic carcinoma cells. Int J Oncol 2014;44:725–34.

[29] Chang YY, Kuo WH, Hung JH, Lee CY, Lee YH, Chang YC, et al. Deregulated microRNAs in triple-negative breast cancer revealed by deep sequencing. Mol Cancer 2015;14:36.

[30] Tang D, Zhang Q, Zhao S, Wang J, Lu K, Song Y, et al. The expression and clinical significance of microRNA-1258 and heparanase in human breast cancer. Clin Biochem 2013;46:926–32.

[31] Liu H, Chen X, Gao W, Jiang G. The expression of heparanase and microRNA-1258 in human non-small cell lung cancer. Tumour Biol 2012;33:1327–34.

[32] Bu P, Wang L, Chen KY, Rakhilin N, Sun J, Closa A, et al. miR-1269 promotes metastasis and forms a positive feedback loop with TGF-beta. Nat Commun 2015;6:6879.

[33] Gonul O, Aydin HH, Kalmis E, Kayalar H, Ozkaya AB, Atay S, et al. Effects of Ganoderma lucidum (Higher Basidiomycetes) Extracts on the miRNA Profile and Telomerase Activity of the MCF-7 Breast Cancer Cell Line. Int J Med Mushrooms 2015;17:231–9.

[34] Emmadi R, Canestrari E, Arbieva ZH, Mu W, Dai Y, Frasor J, et al. Correlative Analysis of miRNA Expression and Oncotype Dx Recurrence Score in Estrogen Receptor Positive Breast Carcinomas. PLoS One 2015;10:e0145346.

[35] Persson H, Kvist A, Rego N, Staaf J, Vallon-Christersson J, Luts L, et al. Identification of new microRNAs in paired normal and tumor breast tissue suggests a dual role for the ERBB2/Her2 gene. Cancer Res 2011;71:78–86.

[36] Lanczky A, Nagy A, Bottai G, Munkacsy G, Szabo A, Santarpia L, et al. miRpower: a web-tool to validate survival-associated miRNAs utilizing expression data from 2178 breast cancer patients. Breast Cancer Res Treat 2016;160:439–46.

[37] Fromm B, Billipp T, Peck LE, Johansen M, Tarver JE, King BL, et al. A Uniform System for the Annotation of Vertebrate microRNA Genes and the Evolution of the Human microRNAome. Annu Rev Genet 2015;49:213–42.

[38] Min P, Li W, Zeng D, Ma Y, Xu D, Zheng W, et al. A single nucleotide variant in microRNA-1269a promotes the occurrence and process of hepatocellular carcinoma by targeting to oncogenes SPATS2L and LRP6. Bull Cancer 2017;104:311–20.

[39] Ma Y, Liang AJ, Fan YP, Huang YR, Zhao XM, Sun Y, et al. Dysregulation and functional roles of miR-183-96-182 cluster in cancer cell proliferation, invasion and metastasis. Oncotarget 2016.

[40] Futreal PA, Coin L, Marshall M, Down T, Hubbard T, Wooster R, et al. A census of human cancer genes. Nat Rev Cancer 2004;4:177–83.

[41] Gosline SJ, Gurtan AM, JnBaptiste CK, Bosson A, Milani P, Dalin S, et al. Elucidating MicroRNA Regulatory Networks Using Transcriptional, Post-transcriptional, and Histone Modification Measurements. Cell Rep 2016;14:310–9.

[42] Woods NK, Padmanabhan J. Inhibition of amyloid precursor protein processing enhances gemcitabine-mediated cytotoxicity in pancreatic cancer cells. J Biol Chem 2013;288:30114–24.

[43] Long J, Ou C, Xia H, Zhu Y, Liu D. MiR-503 inhibited cell proliferation of human breast cancer cells by suppressing CCND1 expression. Tumour Biol 2015;36:8697–702.

[44] Reis-Filho JS, Savage K, Lambros MB, James M, Steele D, Jones RL, et al. Cyclin D1 protein overexpression and CCND1 amplification in breast carcinomas: an immunohistochemical and chromogenic in situ hybridisation analysis. Mod Pathol 2006;19:999–1009.

[45] Werner H, Sarfstein R. Transcriptional and epigenetic control of IGF1R gene expression: implications in metabolism and cancer. Growth Horm IGF Res 2014;24:112–8.

[46] Ogasawara S, Kiyota Y, Chuman Y, Kowata A, Yoshimura F, Tanino K, et al. Novel inhibitors targeting PPM1D phosphatase potently suppress cancer cell proliferation. Bioorg Med Chem 2015;23:6246–9.

[47] Nagy JA, Chang SH, Dvorak AM, Dvorak HF. Why are tumour blood vessels abnormal and why is it important to know? Br J Cancer 2009;100:865–9.

[48] Xu JB, Bao Y, Liu X, Liu Y, Huang S, Wang JC. Defective expression of transforming growth factor beta type II receptor (TGFBR2) in the large cell variant of non-small cell lung carcinoma. Lung Cancer 2007;58:36–43.

[49] Baselga J. Targeting the phosphoinositide-3 (PI3) kinase pathway in breast cancer. Oncologist 2011;16 Suppl 1:12–9.

[50] Chalhoub N, Baker SJ. PTEN and the PI3-kinase pathway in cancer. Annu Rev Pathol 2009;4:127–50.

[51] Waugh MG. Chromosomal Instability and Phosphoinositide Pathway Gene Signatures in Glioblastoma Multiforme. Mol Neurobiol 2016;53:621–30.

[52] Ying Z, Xie X, Chen M, Yi K, Rajagopalan S. Alpha-lipoic acid activates eNOS through activation of PI3-kinase/Akt signaling pathway. Vascul Pharmacol 2015;64:28–35.

[53] Yun YR, Won JE, Jeon E, Lee S, Kang W, Jo H, et al. Fibroblast growth factors: biology, function, and application for tissue regeneration. J Tissue Eng 2010;2010:218142.

[54] Zhu Y, Qiu P, Ji Y. TCGA-assembler: open-source software for retrieving and processing TCGA data. Nat Methods 2014;11:599–600.

[55] Minoche AE, Dohm JC, Himmelbauer H. Evaluation of genomic high-throughput sequencing data generated on Illumina HiSeq and genome analyzer systems. Genome Biol 2011;12:R112.

[56] Yates A, Akanni W, Amode MR, Barrell D, Billis K, Carvalho-Silva D, et al. Ensembl 2016. Nucleic Acids Res 2016;44:D710–6.

[57] Li Y, Qiu C, Tu J, Geng B, Yang J, Jiang T, et al. HMDD v2.0: a database for experimentally supported human microRNA and disease associations. Nucleic Acids Res 2014;42:D1070–4.

[58] Jiang Q, Wang Y, Hao Y, Juan L, Teng M, Zhang X, et al. miR2Disease: a manually curated database for microRNA deregulation in human disease. Nucleic Acids Res 2009;37:D98–104.

[59] Xie B, Ding Q, Han H, Wu D. miRCancer: a microRNA-cancer association database constructed by text mining on literature. Bioinformatics 2013;29:638–44.

[60] Hinske LC, Franca GS, Torres HA, Ohara DT, Lopes-Ramos CM, Heyn J, et al. miRIAD-integrating microRNA inter- and intragenic data. Database (Oxford) 2014;2014.

[61] Gupta S, Dingerdissen H, Ross KE, Hu Y, Wu CH, Mazumder R, et al. DEXTER: Disease-Expression Relation Extraction from Text. Database (Oxford) 2018;2018.

[62] Kibbe WA, Arze C, Felix V, Mitraka E, Bolton E, Fu G, et al. Disease Ontology 2015 update: an expanded and updated database of human diseases for linking biomedical knowledge through disease data. Nucleic Acids Res 2015;43:D1071–8.

[63] Wu TJ, Schriml LM, Chen QR, Colbert M, Crichton DJ, Finney R, et al. Generating a focused view of disease ontology cancer terms for pan-cancer data integration and analysis. Database (Oxford) 2015;2015:bav032.

[64] Chou CH, Chang NW, Shrestha S, Hsu SD, Lin YL, Lee WH, et al. miRTarBase 2016: updates to the experimentally validated miRNA-target interactions database. Nucleic Acids Res 2016;44:D239–47.

[65] Xiao F, Zuo Z, Cai G, Kang S, Gao X, Li T. miRecords: an integrated resource for microRNA-target interactions. Nucleic Acids Res 2009;37:D105–10.

[66] Dweep H, Gretz N. miRWalk2.0: a comprehensive atlas of microRNA-target interactions. Nat Methods 2015;12:697.

[67] Li G, Ross KE, Arighi CN, Peng Y, Wu CH, Vijay-Shanker K. miRTex: A Text Mining System for miRNA-Gene Relation Extraction. PLoS Comput Biol 2015;11:e1004391.

[68] Vergoulis T, Vlachos IS, Alexiou P, Georgakilas G, Maragkakis M, Reczko M, et al. TarBase 6.0: capturing the exponential growth of miRNA targets with experimental support. Nucleic Acids Res 2012;40:D222–9.

[69] Jacobsen A, Silber J, Harinath G, Huse JT, Schultz N, Sander C. Analysis of microRNA-target interactions across diverse cancer types. Nat Struct Mol Biol 2013;20:1325–32.

[70] Consortium GO. Gene Ontology Consortium: going forward. Nucleic Acids Res 2015;43:D1049–56.

[71] Mi H, Poudel S, Muruganujan A, Casagrande JT, Thomas PD. PANTHER version 10: expanded protein families and functions, and analysis tools. Nucleic Acids Res 2016;44:D336–42.

[72] Varet H, Brillet-Gueguen L, Coppee JY, Dillies MA. SARTools: A DESeq2- and EdgeR-Based R Pipeline for Comprehensive Differential Analysis of RNA-Seq Data. PLoS One 2016;11:e0157022.

[73] Dillies MA, Rau A, Aubert J, Hennequet-Antier C, Jeanmougin M, Servant N, et al. A comprehensive evaluation of normalization methods for Illumina high-throughput RNA sequencing data analysis. Brief Bioinform 2013;14:671–83.

[74] Tam S, Tsao MS, McPherson JD. Optimization of miRNA-seq data preprocessing. Brief Bioinform 2015;16:950–63.

[75] Dingerdissen HM, Torcivia-Rodriguez J, Hu Y, Chang TC, Mazumder R, Kahsay R. BioMuta and BioXpress: mutation and expression knowledgebases for cancer biomarker discovery. Nucleic Acids Res 2018;46:D1128–D36.

[76] Consortium U. UniProt: a hub for protein information. Nucleic Acids Res 2015;43:D204–12.

[77] Forbes SA, Bindal N, Bamford S, Cole C, Kok CY, Beare D, et al. COSMIC: mining complete cancer genomes in the Catalogue of Somatic Mutations in Cancer. Nucleic Acids Res 2011;39:D945–50.

[78] Szklarczyk D, Franceschini A, Wyder S, Forslund K, Heller D, Huerta-Cepas J, et al. STRING v10: protein-protein interaction networks, integrated over the tree of life. Nucleic Acids Res 2015;43:D447–52.

